# Integrative Omics Identifies Candidate Plasma Biomarkers and Cellular Targets Associated with Thoracic Aortic Aneurysm

**DOI:** 10.1101/2025.07.25.666909

**Authors:** Alexander C. Bashore, Emir Radkevich, Vladimir Roudko, Zhihong Chen, Kai Nie, Tia Rizakos, Jarod Morgenroth-Rebin, Darwin D’Souza, Anastasia Filimonov, Abena Gyasi, Abrisham Eskandari, Mariya Shadrina, Ana Devesa, Enrique J. Garcia, Maria Trivieri, Georgios Soultanidis, Shams Rashid, Matthew Tomey, Stamatios Lerakis, Barbara A. Sampson, Sikander Hayat, Seunghee Kim-Schulze, Ismail El-Hamamsy, Valentin Fuster, Amy R. Kontorovich

## Abstract

**Objectives:** To define the cellular landscape of thoracic aortic aneurysm (TAA) and identify circulating biomarkers associated with disease burden.

**Background:** TAA is marked by progressive aortic wall weakening and dilation, predisposing to rupture and dissection. However, its cellular architecture remains incompletely defined, and reliable circulating biomarkers are lacking.

**Methods:** Single-cell RNA sequencing was performed on 17 aortic tissue samples from 10 patients undergoing TAA repair and integrated with publicly available datasets to characterize disease-associated cell states. In parallel, tomographic imaging and plasma proteomics were used to identify biomarkers associated with aortic diameter. Key findings were further assessed through integration with single-cell data, external validation, and in vitro stimulation of primary human adventitial fibroblasts with fibroblast growth factor 23 (FGF-23).

**Results:** We identified 25 cellular subsets, including macrophages, endothelial cells, vascular smooth muscle cells, and fibroblasts, with substantial heterogeneity in cellular composition and transcriptional state. Genome-wide association study candidate genes, including *JUN* and *TPM3*, showed cell type-specific upregulation. Plasma proteomics identified multiple biomarkers associated with aortic diameter, of which FGF-23 was independently validated in the UK Biobank as elevated in individuals with TAA. FGFR1, the receptor for FGF-23, was selectively expressed in fibroblasts and subsets of vascular smooth muscle cells, with strongest downstream signaling in fibroblasts. FGF-23 stimulation induced inflammatory and extracellular matrix remodeling programs in primary human adventitial fibroblasts.

**Conclusions:** These findings define the cellular landscape of TAA and identify the FGF-23-FGFR1 axis as a biomarker-linked pathway that may contribute to aneurysm progression.

**Condensed Abstract:** Thoracic aortic aneurysm (TAA) is characterized by progressive aortic dilation and risk of rupture, yet its cellular architecture and circulating biomarkers remain incompletely defined. We performed single-cell RNA sequencing on 17 aortic samples from 10 patients and integrated these data with public datasets to define the cellular landscape of TAA. In parallel, imaging and plasma proteomics (n=10) were used to identify biomarkers associated with aortic diameter. We identified 25 cellular subsets, including macrophages, endothelial cells, vascular smooth muscle cells, and fibroblasts, with notable transcriptional heterogeneity and cell type-specific upregulation of GWAS-associated genes (e.g., JUN, TPM3). Plasma proteomics identified fibroblast growth factor 23 (FGF-23) as associated with aortic diameter and elevated in TAA in the UK Biobank. FGFR1, its receptor, was selectively expressed in fibroblasts and VSMCs, and FGF-23 stimulation induced inflammatory and extracellular matrix remodeling programs in fibroblasts. These findings link a circulating biomarker to stromal cell signaling in TAA.

**Highlights:**

- Integrative single-cell RNA sequencing defined a diverse cellular landscape in thoracic aortic aneurysm tissue, identifying 25 distinct cell populations.
- Cross-dataset harmonization revealed marked differences in cellular composition across studies while supporting shared stromal and immune programs in thoracic aortic aneurysm.
- Plasma proteomics identified FGF-23 as a candidate biomarker associated with aortic diameter and independently linked to thoracic aortic aneurysm presence in the UK Biobank
- FGFR1 was enriched in fibroblasts and subsets of vascular smooth muscle cells, and FGF-23 induced inflammatory and extracellular matrix remodeling programs in human primary adventitial fibroblasts.

## Introduction

Thoracic aortic aneurysm (TAA) is a life-threatening condition characterized by progressive enlargement of the thoracic aorta, which can ultimately lead to severe complications such as aortic dissection or rupture if left untreated. The incidence of TAA is estimated to be approximately 10 per 100,000 person-years^1^. However, the growing availability of advanced imaging modalities and the increasing age of the general population, both of which have contributed to a higher detection rate of previously undiagnosed cases^2,3^, suggest that this is likely an underestimate. Aortic dissection, an often fatal complication of TAA, occurs when a tear in the aortic wall allows blood to flow between its layers, weakening its structural integrity and compromising perfusion. Currently, the primary approach to mitigating the risk of aortic dissection is careful monitoring of aneurysm growth through regular imaging, and elective surgical intervention when the aneurysm reaches a diameter that indicates a high risk for rupture or dissection^4–6^.

TAA pathobiology involves weakening of the aortic wall^7^, driven by processes like extracellular matrix (ECM) degradation^8^, elastin fragmentation^9^, and vascular smooth muscle cell (VSMC) proliferation^10^ and phenotypic switching^11–13^. Despite these well-established pathological mechanisms, the full range of molecular and cellular processes leading to aortic degradation and TAA formation remain poorly understood. Consequently, recent efforts have focused on utilizing rapidly advancing single-cell technologies to unravel the complete cellular landscape within TAA tissue and to identify causal cell types and molecular drivers of disease progression^14–18^. However, these studies have produced conflicting results, with one suggesting that it is a T cell mediated inflammatory disease^16^ while another indicates the disease process is characterized by alterations in VSMC and fibroblasts^15^. These discrepancies make it challenging to identify causal regulators of the disease process and better understand the underlying disease mechanisms. Therefore, a comprehensive description of the cellular processes mediating TAA formation and subsequent dissection is crucial for identifying molecular regulators of the disease and facilitating the development of novel therapeutics.

To address some of these discrepancies, and to identify circulating plasma biomarkers that reflect aneurysm presence and disease burden, we generated a transcriptional dataset of TAA tissue using single-cell RNA-sequencing (scRNA-seq) on a large population of cells derived from aneurysmal tissue. TAA tissue was characterized by substantial cellular heterogeneity as well as previously undescribed subtypes of endothelial cells (ECs), VSMCs, and fibroblasts. By integrating these cell-type-resolved transcriptional profiles with plasma proteomics, we identified circulating factors that map to specific stromal signaling programs and may contribute to TAA pathobiology.

## Methods

### Human Studies

All human subjects research in this study, including the use of human tissues, conformed to the principles outlined in the Declaration of Helsinki. All patient information was de-identified. For single-cell studies, tissues from human ascending aorta were collected from ten patients at time of TAA surgery. These human subject studies were performed with approval (STUDY-21-01302) of the local Institutional Review Board (IRB) of the Icahn School of Medicine at Mount Sinai (ISMMS), and written informed consent was obtained from all participants. Exclusion criteria included current pregnancy, age < 18 years and cognitive impairment. **Table 1** summarizes the demographic and clinical characteristics of the entire cohort and patient-level data is in **Table S1**.

**Table.** Summary of Patient Characteristics for Cohort used in study.

|  |  |
| --- | --- |
| **N** | **11** |
| **Age (median)** | **59** |
| **Male sex** | **10 (91%)** |
| **Maximal dilatation at SoV** | **3 (27%)** |
| **Maximal dilatation at ascending segment** | **8 (73%)** |
| **Maximal aneurysmal diameter (median)** | **5.1** |
| **Dissection present** | **1 (9%)** |
| **Any smoking history** | **4 (36%)** |
| **Diabetes** | **1 (9%)** |
| **Hypertension** | **6 (55%)** |
| **COPD** | **0 (0%)** |
| **AVR** | **2 (18%)** |
| **Reoperation** | **0 (0%)** |
| **Bicuspid Aortic Valve** | **3 (27%)** |
| **Atherosclerosis** | **5 (46%)** |
| **FBN1+** | **1 (9%)** |
| **TGFBR1+** | **1 (9%)** |
| **Marfan Syndrom** | **1 (9%)** |
| **Loeys-Dietz Syndrome** | **1 (9%)** |
| **scRNA-seq** | **10 (91%)** |
| **Olink** | **10 (91%)** |

### Imaging

Participants underwent tomographic imaging of the thoracic aorta prior to TAA surgery. Modalities included cardiac magnetic resonance imaging (CMR), magnetic resonance angiography (MRA) or gated contrast tomography angiography (CTA). Thoracic aortic dimensions were measured and reported via established standards per modality.

### Genomic sequencing

Participants underwent either clinical genetic testing on a targeted gene panel for aortopathy and/or whole exome sequencing (Azenta Life Sciences, Illumina platform, Human Twist capture kit). For the latter, vcf files were created from raw fastq files and joint variant detection was performed for cases using GATK (v3.8). Analysis was restricted to capture regions pertaining to 23 genes of interest (COL3A1,FBN1, SMAD3, TGFB2, TGFBR1, TGFBR2, ACTA2, MYH11, MYLK, LOX, PRKG1, EFEMP2, ELN, FBN2, FLNA, NOTCH1, SLC2A10, SMAD4, SKI, CBS, COL4A4, PKD1, PKD2) based on those previously validated for clinical association with TAA as per the ClinGen expert working group^19^. Resulting VCF files were annotated using Annovar(4) and applying the following filters using in-house scripts: 1) B-allele frequency (BAF) = 0.3-0.7 for heterozygous calls; 2) BAF >= 0.9 for homozygous calls; genotype quality >= 30; read depth >= 7. Putatively deleterious variants were defined as those that were either previously classified as likely pathogenic or pathogenic in the ClinVar database by at least one Badge laboratory. Possibly deleterious variants were defined as rare (MAF < 0.1% in the gnomAD database) and without likely benign or benign classification by any Badge laboratory in ClinVar.

### Aortic tissue cell dissociation for single-cell RNA-seq

Freshly dissected transmural samples (1-3 per anatomic segment per participant) were collected from the aneurysmal section of the aorta, the samples were then transported from the pathology department to the laboratory in tissue storage buffer (Miltenyi, 130-100-007) within 2 hours of removal. Upon arrival, the tissues were placed in a glass Petri dish, and any remaining connective tissue was carefully removed. The tissue was then weighed and minced with a scalpel into small pieces of approximately 1 mm³ in size. To generate a single-cell suspension, a Multi-Tissue Dissociation Kit 2 (Miltenyi, 130-110-203) was used. Briefly, the minced tissue pieces were transferred to a gentleMACS C Tube (Miltenyi, 130-096-334) containing the enzymatic mix. The tubes were then placed in a gentleMACS Octo Dissociator (Miltenyi), and the pre-programmed Multi_G dissociation program was executed. Following dissociation, the tissue was filtered through a 70 µm strainer, washed with 3 mL of RPMI + 20% FBS, and centrifuged at 600 x g for 5 minutes. The resulting pellet was resuspended in PBS + 0.05% BSA for subsequent single-cell RNA sequencing analysis.

### scRNA-seq

Viability of single cells was assessed using Acridine Orange/Propidium Iodide viability staining reagent (Nexcelom), and debris-free suspensions of >80% viability were deemed suitable for the experiments. ScRNA-Seq was performed using the Chromium platform (10x Genomics) with the 5’ gene expression (5’ GEX) V2 kit, with a targeted recovery of 10,000 cells. Briefly, Gel-Bead in Emulsions (GEMs) were generated on the sample chip in the Chromium X system. Barcoded cDNA was extracted from the GEMs after Post-GEM RT-cleanup and amplified for 12 cycles. Full-length cDNA from poly-A mRNA transcripts underwent enzymatic fragmentation and size selection to optimize cDNA amplicon size (∼400 bp) for library construction. The cDNA fragments were then subjected to end-repair, adapter ligation, and 10x-specific sample indexing according to the manufacturer’s protocol (10x Genomics). The concentration of the single-cell library was accurately quantified using qPCR (Kapa Biosystems) to achieve appropriate cluster counts for paired-end sequencing on the NovaSeq 6000 (Illumina), aiming for a sequencing depth of 25,000 reads per cell. Raw FASTQ files were aligned to the GRCh38 reference genome (2020-A) using Cell Ranger v5.0.1 (10x Genomics).

### Single-cell RNA-seq analysis

FASTQ files for Gene Expression (GEX) libraries were processed using the Cellranger Count pipeline (version 5.0.1) with the GRCh38-2020-A reference genome provided by 10x Genomics. This pipeline performed filtering, barcode counting, UMI counting, and cell identification. Samples of low quality were excluded, defined as those with fewer than 500 median genes per cell and fewer than 2,000 median UMIs per cell. Following sample level filtering, low-quality cells, defined as those with fewer than 200 detected genes or more than 25,000 total gene counts, as well as dying cells exhibiting greater than 15% mitochondrial gene content, were excluded from downstream analysis. Quality control metrics for each sample are detailed in **Table S2**.

The obtained expression data were normalized, log-transformed, and projected into a 2D latent space using the Scanpy pipeline (v.1.10.2) as previously described^20^. Principal component analysis (PCA) was performed, and the top 50 principal components were used to calculate uniform manifold approximation and projection (UMAP) coordinates. The Leiden algorithm was applied to construct a shared nearest-neighbor graph for clustering. Cluster marker genes were identified using a *t*-test with overestimated variance, following the “one-versus-everyone” strategy implemented in Scanpy. The top 100 upregulated and downregulated genes for each cluster are detailed in **Table S3**. Clusters enriched for marker genes corresponding to two or more distinct cell phenotypes were classified as doublets and subsequently removed from the analysis.

### Reference-based mapping of human aneurysm public datasets

Two external datasets^15,16^ were downloaded and processed using the same pipeline as described earlier. Reference-based mapping was performed, and the external datasets were projected onto our annotated dataset using the SingleR method, following the developers’ recommended guidelines.

### Compositional analysis

To investigate compositional changes across cell types, we performed differential composition analysis using the scCODA (Bayesian modeling of compositional single-cell data) framework implemented in the pertpy package v.0.10.0^21^. This method accounts for the compositional nature of cell-type proportions in single-cell datasets, where the relative, rather than absolute, abundances are more informative. All the analysis was implemented using default parameters, including automatic reference cell type selection during model preparation.

### Differential gene expression between aneurysm and control samples

We employed decoupler^22^ to generate pseudobulk profiles prior to conducting differential gene expression analysis. Each cell type was analyzed independently, with rigorous quality control applied to ensure robust results. Genes expressed in fewer than 5% of cells within each cell type were excluded, along with cells expressing fewer than 200 genes. Differential gene expression between conditions was assessed using the pseudobulk approach implemented in pydeseq2 (v.0.5.0)^23^, which provides a statistically robust framework for comparing gene expression across aggregated samples. This methodology enhances power and mitigates the noise inherent to single-cell data, allowing for reliable detection of condition-specific transcriptional changes.

### Analysis of public mouse aneurysm data

Publicly available mouse scRNA-seq data from an angiotensin II-induced aneurysm model were obtained^24^. The data were processed and analyzed following the same pipeline used for our scRNA-seq analysis.

### Proteomic assessment

Whole blood was collected immediately prior to surgery via venipuncture into a Vacutainer containing EDTA as an anticoagulant and processed by the Human Immune Monitoring Core of the ISMMS. The blood sample was then centrifuged at 2,000 x g for 10 minutes at 4°C to separate the plasma. The resulting plasma was carefully aliquoted and stored at -80°C until ready for analysis. Proteomic profiling was performed on plasma samples using the Olink Target 96 Inflammation panel (Olink Proteomics), which quantifies 92 inflammation-related proteins using the Proximity Extension Assay (PEA) technology. A mixture of oligonucleotide-labeled antibody pairs targeting each protein was introduced into the samples and left to incubate at 4°C for 16 hours. Two distinct epitope-specific antibodies were used for each protein. The presence of the target protein in the sample brought the partner probes in close proximity, leading to the formation of a double-strand oligonucleotide serving as a template for polymerase chain reaction (PCR) amplification. Subsequently, the extension master mix was introduced to the sample, initiating the detection of specific target sequences and the generation of amplicons using PCR in 96-well plates. For protein detection, a Dynamic array Integrated Fluidic Circuit (IFC) 96 × 96 chip was primed, loaded with 92 protein-specific primers, and combined with sample amplicons. Real-time microfluidic quantitative PCR (qPCR) was carried out using the Biomark system (Fluidigm) to quantify the target proteins. Data are presented as Normalized Protein Expression (NPX) values, a relative quantification metric derived from Olink’s proximity extension assay (PEA). NPX values are log2-transformed and normalized to minimize technical variation, enabling robust comparison of protein abundance across samples within the same assay.

### Candidate Biomarker associations with TAA diameter

To assess the relationship between each plasma analyte and TAA diameter, a ROUT test (Q = 5%) was first applied to identify and remove outliers, ensuring data quality. Following this, a correlation matrix was generated to evaluate the associations between plasma analytes and aortic diameter. A simple linear regression was then performed for candidate biomarkers of interest. Key associations of interest were subsequently visualized using scatter plots to illustrate the strength and direction of the relationships.

### Analysis of UK Biobank Olink data

Summary statistics from the UK Biobank were accessed via the interactive web tool (https://proteome-phenome-atlas.com/), developed as part of a recent study that profiled 2,923 unique plasma proteins using Olink Explore™ Proximity Extension Assay and next-generation sequencing in 53,026 adults^25^. Incident disease-wide association results for “Thoracic Aortic Aneurysm” were queried across the full population, applying a significance threshold of *p* < 0.05. The analysis identified 357 proteins with positive associations and 94 proteins with negative associations among 166 cases (defined as individuals with ICD-10 codes I71.0, I71.1, and I71.2) and 50347 controls without these ICD-10 codes.

### STRING Analysis

Network analysis was performed using stringAPP v2.1.1 in Cytoscape v3.8. The protein query function for FGF-23 identified potential interacting partners based on known and predicted protein-protein associations, integrating data from experimental evidence and curated databases.

### FGFR1-Associated Gene Module Scoring

FGFR1-associated transcriptional module scores were calculated using the score_genes function in Scanpy^20^. For each cell, the module score was defined as the average expression of genes within a predefined FGFR1-associated gene set, normalized by subtracting the average expression of a set of control genes matched for overall expression levels. Control genes were selected by binning all genes according to their mean expression and randomly sampling from each bin to account for technical and abundance-related biases. Default parameters were used for all analyses.

### Bulk RNA Sequencing of Primary Human Adventitial Fibroblasts

Primary human adventitial fibroblasts (Lonza: CC-7014) were cultured in stromal cell growth medium (Lonza: CC-3205) according to the manufacturer’s instructions. Cells were passaged to P3, and 7.5 × 10 cells were seeded per well in 12-well plates and allowed to adhere for 24 hours. Cells were then treated with recombinant human FGF-23 (25 ng/mL) (Thermo Fisher Scientific: 100-52) for 48 hours, with vehicle-treated cells serving as controls (0.1% bovine serum albumin).

Total RNA was isolated using the Quick-RNA MiniPrep Kit (Zymo Research: R1055), and RNA integrity was assessed prior to library preparation. RNA samples were quantified using a Qubit fluorometer and RNA integrity was evaluated using an Agilent TapeStation. RNA sequencing libraries were prepared using the NEBNext Ultra II RNA Library Prep Kit for Illumina with poly(A) mRNA enrichment. Briefly, mRNA was isolated using oligo(dT) beads, fragmented, and reverse transcribed to generate first- and second-strand cDNA, followed by end repair, adapter ligation, and PCR amplification. Libraries were validated and quantified prior to sequencing. Sequencing libraries were multiplexed and sequenced on an Illumina NovaSeq platform using a 2 × 150 bp paired-end configuration. Raw base call files were converted to FASTQ format and demultiplexed using bcl2fastq (Illumina). Reads were trimmed for adapter sequences and low-quality bases using Trimmomatic (v0.36) and aligned to the human reference genome (GRCh38/hg38) using STAR (v2.5.2b). Gene-level counts were generated using feature Counts (Subread v1.5.2), considering uniquely mapped reads overlapping exon regions.

Raw RNAseq counts were processed in R (4.5.2) using DESeq2^26^. Briefly, genes with < 10 counts in < 3 samples were excluded prior to analysis. Sample quality was assessed using 4 orthogonal metrics: 1) library size deviation from the median (> 50% deviation considered an outlier), 2) Mahalanobis distance in PCA space of variance stabilized counts (VST) (distances exceeding 97.5^th^ percentile of the χ² distribution wwas considered an outlier, p<0.025 with df (degrees of freedom)=2)^27^, 3) hierarchical clustering of inter sample Euclidean distances, and 4) mean inter sample Pearson correlation (< 0.95 threshold)^28^. Any sample identified as an outlier by ≥ 3 metrics was excluded from the analysis. Gene-level outliers were identified via Cook’s distance^29^ with affected genes automatically excluded from multiple testing correction by DESeq2. Differential expression (DE) was performed with the Wald test and log2 fold change values were shrunken using “apeglm” method^30^. Multiple testing correction was performed with DESeq2’s native Benjamini-Hochberg procedure with independent filtering^31^. DE genes were defined by FDR <0.05 and |log2FC|>0.26 (≥1.2-fold change).

### Code for the analysis

Python code used in the single-cell/single-nuclei data analysis is deposited in Github, as a jupyter notebook [https://github.com/ismms-himc/Bashore_aneurysm].

## Data and Materials Availability

All data will be released on the Gene Expression Omnibus (GEO) upon acceptance of this article.

## Results

### scRNA-seq reveals 25 distinct cell types in TAA tissue

To investigate the cellular diversity within TAA tissue, we obtained aneurysmal aortic tissue from patients undergoing corrective surgery. Baseline characteristics of the patients are summarized in **Table 1 and Table S1**. Enzymatic digestion was performed on 17 tissue samples obtained from 10 participants, generating cell suspensions for scRNA-seq. Additionally, 11 patients, including 9 of the 10 patients that also underwent scRNA-seq, completed pre-surgical imaging and blood collection for subsequent plasma proteomics analysis (**Figure 1A**). After quality control and filtering (**Table S2**), our scRNA-seq analysis incorporated data from 98,983 cells across all samples. Visualization using UMAP revealed 25 distinct cell populations (**Figure 1B**). Interestingly, clusters 4 and 5 were composed almost entirely of cells from patients 7001 and 7005, respectively; due to this strong donor bias, these clusters were excluded from subsequent analyses to avoid confounding effects related to individual patient variability rather than disease-specific biological processes. Using established canonical markers, we identified multiple immune and stromal cell populations, including 3 macrophages, 1 monocyte, 1 dendritic cell, 5 T cells, 1 B cell, 1 NK cell, 1 mast cell, 4 ECs, 4 VSMCs, 2 fibroblasts, and 1 mesothelial cell populations (**Figure 1C**).

**Figure 1.**
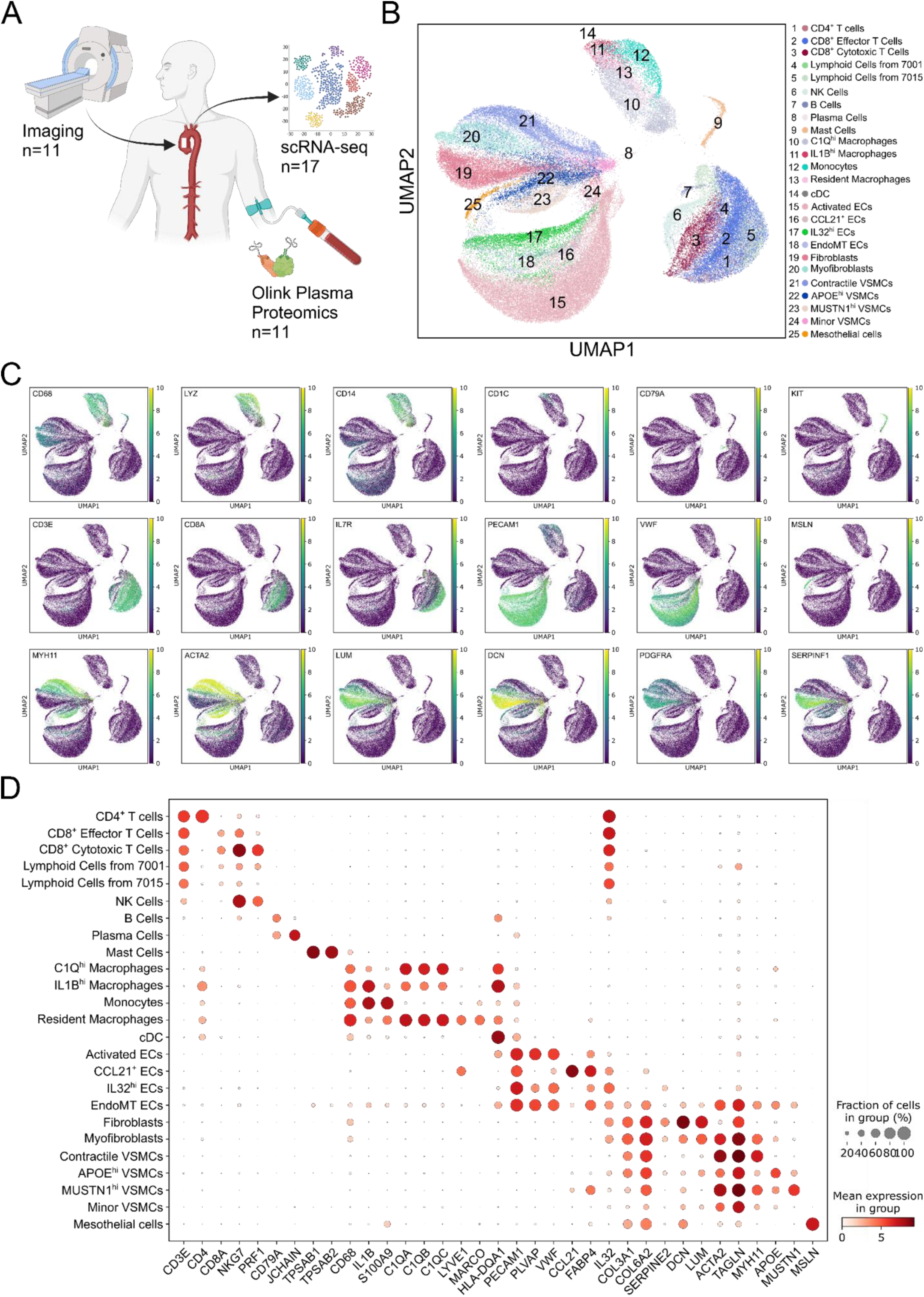
Single-cell RNA-seq analysis of aneurysmal tissue reveals 26 distinct cell populations. (A) Schematic of the experimental workflow integrating imaging, plasma proteomics, and single-cell RNA sequencing (scRNA-seq) for comprehensive analysis of aneurysmal tissue. For scRNA-seq, 17 samples were acquired from 10 patients. (B) UMAP visualization of scRNA-seq data, showing 25 distinct cell clusters. (C) Feature plots illustrating the expression of key marker genes across various cell types: macrophages (CD68, LYZ, CD14), dendritic cells (CD1C), B cells (CD79A), mast cells (KIT), T cells (CD3E, CD8A, IL7R), endothelial cells (PECAM1, VWF), mesothelial cells (MSLN), vascular smooth muscle cells (VSMCs; MYH11, ACTA2), myofibroblasts (LUM, DCN), and fibroblasts (PDGFRA, SERPINF1). (D) Dot plot displaying the select marker genes for each identified cell cluster used to annotate subsets of similar cell types.

To characterize the identified cell types in our dataset further, we analyzed the select top marker genes for each cluster (**Figure 1D and Table S3**). Our analysis revealed intriguing patterns, such as macrophages enriched in IL1B (Cluster 11) and C1Q^hi^ macrophages (Cluster 10), that have been described in other disease contexts^32,33^ but not previously in TAA. It also revealed an unprecedented diversity among ECs, including classical activated ECs (Cluster 15). In addition, fibrotic ECs (Cluster 18), characterized by the expression of *ACTA2*, *TAGLN*, and *MYH11*, indicating they are going through endothelial-to-mesenchymal transition (EndoMT), as well as distinct EC subsets expressing *IL32* (Cluster 17) and *CCL21* (Cluster 16), were identified. The dataset also revealed a wide variety of VSMCs, including contractile VSMCs enriched in *ACTA2* and *MYH11* (Cluster 21), as well as APOE^hi^ (Cluster 22) and MUSTN1^+^ (Cluster 21) VSMCs, suggesting roles in lipid metabolism and ECM remodeling, respectively. Our analysis of fibroblasts identified two distinct subsets: quiescent fibroblasts (cluster 19) and myofibroblasts (cluster 20). The myofibroblast subset expressed both VSMC and fibroblast genes, suggesting it may originate from both cellular lineages. Several of these cell types have not been described previously. These findings highlight the cellular complexity within TAA tissue, providing novel insights into the diverse cell populations contributing to aneurysm pathology.

### Comparison across datasets of cellular alterations in TAA tissue

Given the discrepancies between previously published studies, we reanalyzed the datasets from Li *et al.*^16^ and Chou *et al.*^15^. Variations in single-cell analysis outcomes and conclusions can arise from multiple factors, including the timing of tissue acquisition, methods of tissue dissociation and processing, and the analytical approaches used. Since the datasets have already been generated, the only variable we could standardize was the analysis method. To address this, we reanalyzed these publicly available datasets using the same analytical pipeline. We first aligned each dataset to our scRNA-seq reference to ensure consistent annotation of cell types. When visualizing the Li (**Figure 2A**) and Chou (**Figure 2E**) datasets, we identified all the cell types described in our own dataset. However, notable differences emerged: the Li dataset showed an overrepresentation of immune cells, particularly T cells within the aneurysm group (**Figure 2B and C**), while the Chou dataset was dominated by contractile VSMCs (**Figure 2F and G**). These findings align with what was reported in their respective publications, highlighting differences in cell type proportions across studies. To verify these observations, we conducted a compositional analysis using scCODA^34^, which revealed an expansion of CD4^+^ T cells and CD8^+^ effector T cells, alongside a reduction in contractile VSMCs in aneurysmal tissue from the Li study (**Figure 2D**). In contrast, the Chou study identified a decrease in resident macrophages and fibroblasts, with an increase in contractile VSMCs in aneurysmal tissue (**Figure 2H**). This compositional analysis underscores the striking differences in tissue composition between the two studies.

**Figure 2.**
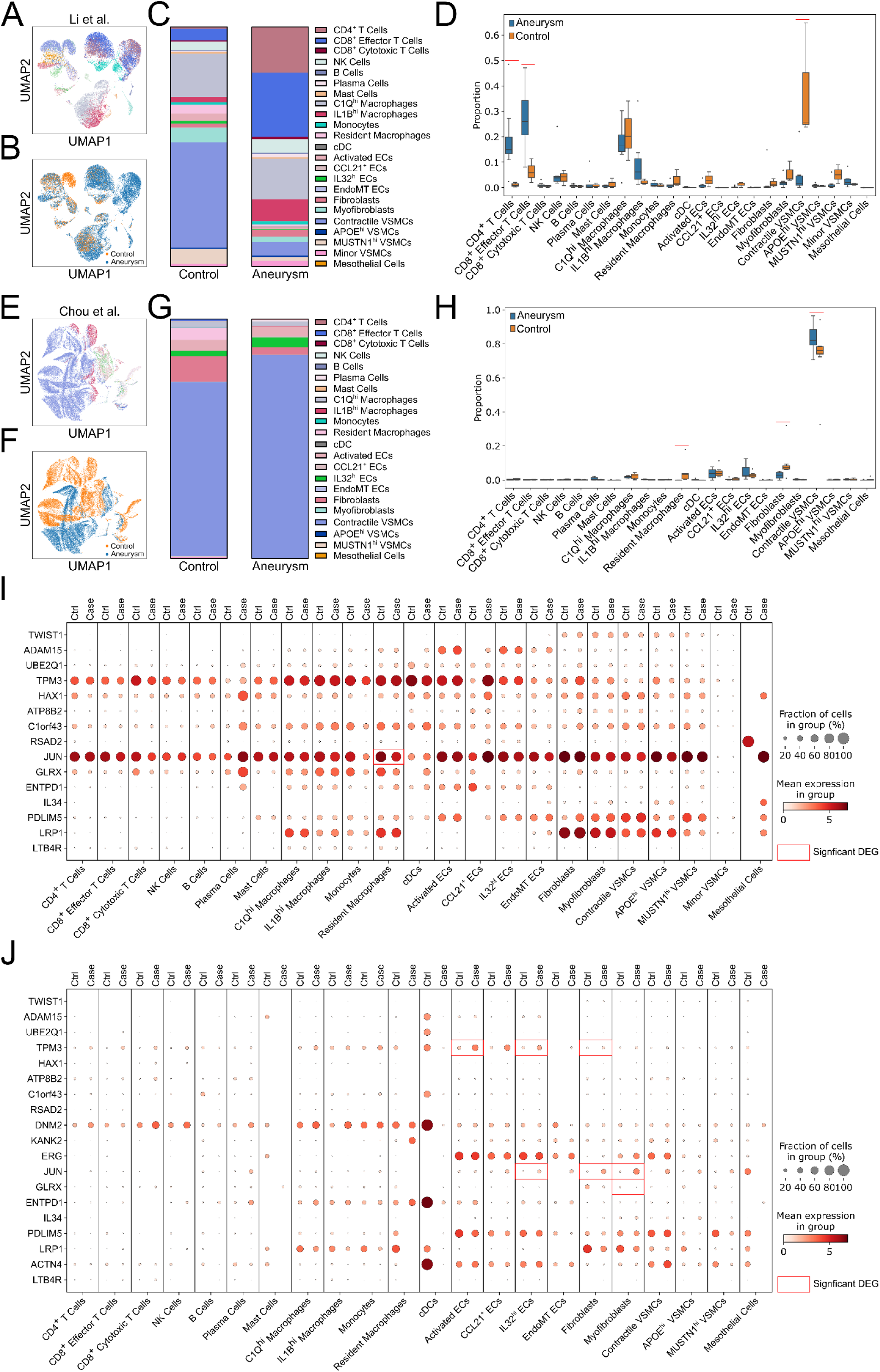
Comparative analysis of publicly available datasets. (A and B) UMAP visualization of Li et al. (A and B) and Chou et al. (E and) datasets, identifying all cellular subsets present in our dataset. (C and G) Stacked bar graph comparing proportions of each cellular subset between control and aneurysmal tissue in respective studies. (D and H) scCODA analysis revealing significant shifts in cell type composition between control and aneurysmal tissue across the two independent datasets. (I andJ) Dot plots depicting the expression of differentially expressed (DE) genes prioritized as targets of aneurysm-associated SNPs, shown within the Li et al. dataset (I) and the Chou et al. dataset (J).

Both studies reported widespread gene expression differences and identified differentially expressed (DE) genes linked to potential GWAS targets. We reanalyzed their data within our pipeline using a pseudobulk approach, which reduces the likelihood of false discoveries^35^. We then assessed the expression of these GWAS target genes in both datasets (**Figure 2I and J**) to determine whether any were differentially expressed in aneurysmal tissue compared to control tissue. In the Li et al. dataset, the only differentially expressed gene was *JUN* (**Figure 2I**), which was downregulated in resident macrophages in aneurysmal tissue. In contrast, several genes exhibited differential expression in the Chou et al. dataset (**Figure 2J**). Specifically, *TPM3* was upregulated in activated ECs, IL32^hi^ ECs, and fibroblasts in aneurysmal tissue; *JUN* was elevated in IL32^hi^ ECs, fibroblasts, and myofibroblasts; and *GLRX* was downregulated in myofibroblasts. Interestingly, we did not detect any changes in *LRP1* expression within any cellular subset, a gene that was emphasized as of potential interest in the Chou et al. study. It is important to note that while the Li et al. study employed scRNA-seq, the Chou et al. study utilized single-nucleus RNA-seq (snRNA-seq). This technical difference—profiling nuclear RNA rather than whole-cell RNA—could introduce discrepancies between the two studies.

### Cell type specific target genes across studies

To integrate our findings with the Li and Chou studies, we examined the expression of all DE genes associated with GWAS findings from their studies across our dataset (**Figure 3**). The goal was to identify whether any of the proposed critical target genes were uniquely associated with specific cell types in our refined cellular annotation of TAA tissue. Among the prioritized genes from the Li *et al.* study, *ADAM15* was specifically expressed in ECs, with the highest expression in IL32^+^ and activated EC subsets. Similarly, *ERG*, a gene implicated in maintaining normal aortic wall function^36^ and associated with aneurysm^37^, was also enriched in ECs. *KANK* displayed high expression in myofibroblasts, suggesting it could be involved in the development of fibroblasts and VSMCs into this phenotype. Other genes from this study were either not highly expressed or lacked specificity to any one cell population. For the key genes highlighted in the Chou *et al.* study, *IL34* was uniquely expressed in myofibroblasts. Interestingly, *LRP1* showed the highest expression in fibroblasts and myofibroblasts, suggesting these cell populations may mediate its disease-associated effects.

**Figure 3.**
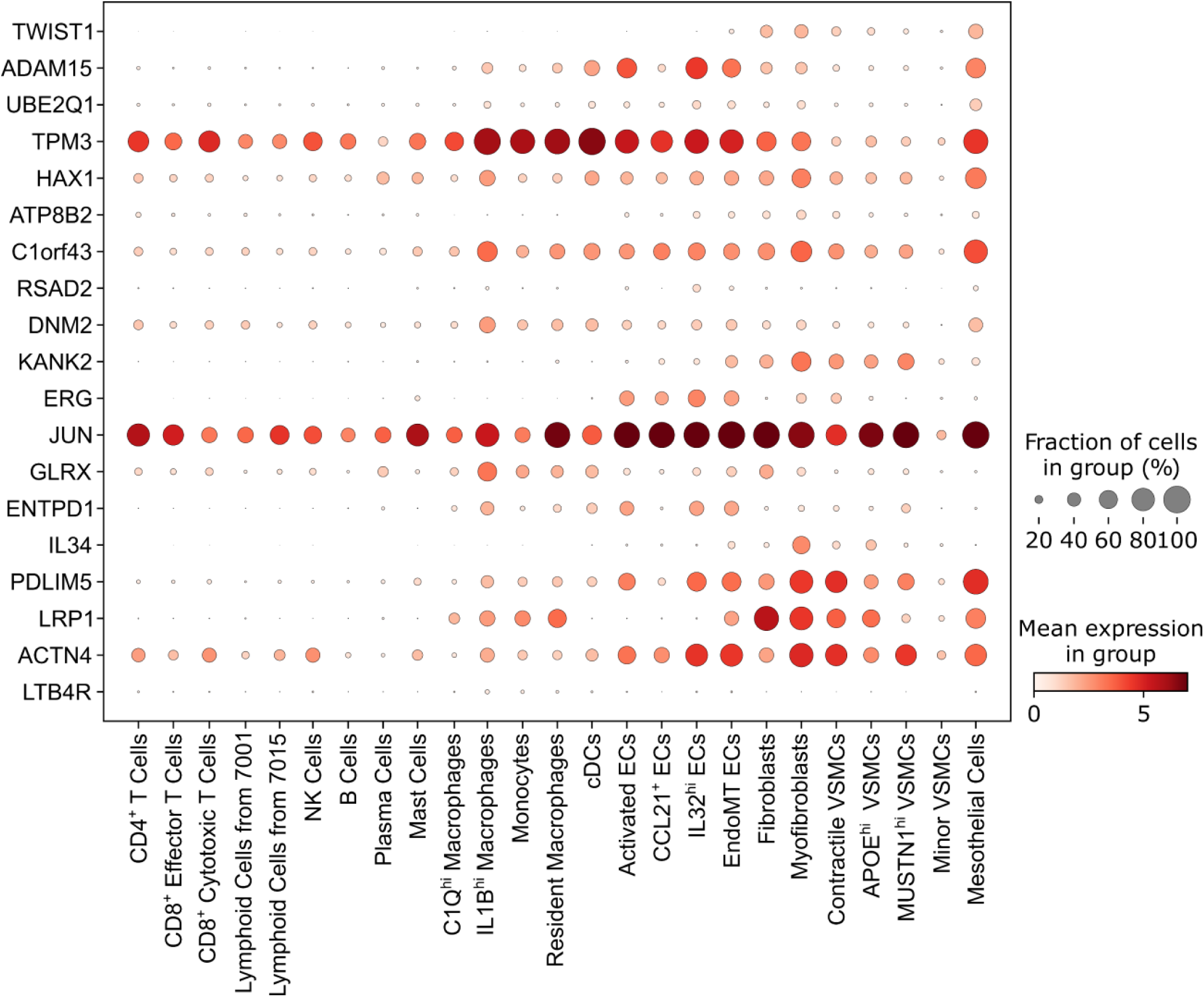
Comparative cell-type enrichment analysis of critical DE genes across three studies. Dot plot visualizing the cell-type-specific expression of GWAS associated DE genes identified in the Li et al. study and Chou et al. study within our dataset.

### FGF-23 is Associated with TAA Burden and Presence

To extend our findings and identify candidate circulating plasma biomarkers reflecting TAA presence and disease burden, we performed a proteomic analysis using a panel of 92 inflammatory analytes (**Table S4**). We began by conducting a correlation analysis between aortic diameter and each analyte in the panel (**Figure 4A**). Our analysis identified several marginal associations with candidate biomarkers such as ADA, HGF, CASP-8, IL8, CCL11, NT-3, FGF-23, TNF, TWEAK, and Flt3L. However, none of these analytes reached the Bonferroni-corrected significance threshold (p<7.04×10⁻), likely due to the limited sample size. To expand our analysis, we leveraged data from a recent study that created a plasma proteome atlas within the UK Biobank^25^. This resource encompasses approximately 3,000 proteins measured in over 50,000 participants across a diverse range of health and disease conditions, offering a valuable tool for deeper investigation. From this atlas, we curated a cohort of 166 participants with TAA (identified by ICD-10 codes I71.0, I71.1, and I71.2) and 50,347 control individuals without these codes. In this dataset, 451 proteins were found to be associated with TAA at a significance level of p<0.05, with 357 showing positive associations and 94 showing negative associations (**Figure 4B**). **Figure 4C** illustrates the top ten positively and top ten negatively associated proteins. Notably, those with the strongest positive associations with TAA include proteins involved in cardiovascular remodeling (NTproBNP, IGFBPs, RNASE1)^38–40^, angiogenesis (PGF, LTBP2)^41,42^, and extracellular matrix remodeling and fibrosis (MMP12, SUSD5)^43,44^. It is unsurprising that none of the proteins identified in our correlation analysis overlap with the top 10 most positively associated proteins in the UK Biobank dataset. This discrepancy likely arises from the UK Biobank’s significantly larger sample size, broader protein coverage, and its case-control design, in contrast to our study, which focused on correlations with aortic size measurements and did not have control participants. Nonetheless, we explored the associations of proteins identified as having a marginal positive correlation with aortic diameter. Interestingly, FGF-23 was the only plasma analyte that demonstrated a significant positive association with TAA in the UK Biobank dataset (**Figure 4D**), supporting its role as a biomarker of aneurysm presence.

**Figure 4.**
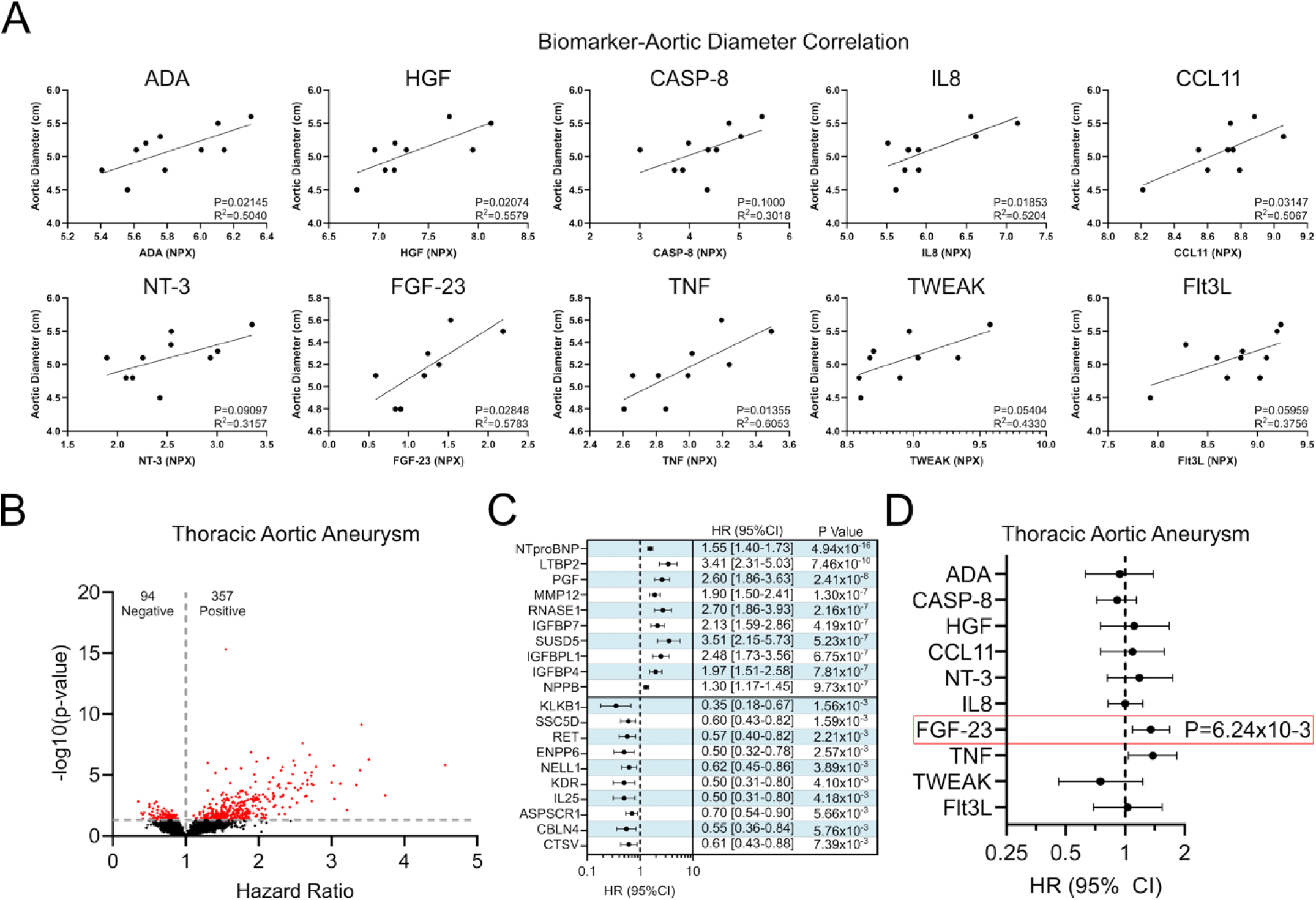
Plasma biomarkers associated with TAA severity. (A) Scatter plots showing the relationship between plasma levels of select biomarkers (measure in NPX) and aortic diameter (cm). Individual biomarkers (ADA, HGF, CASP-8, IL8, CCL11, NT-3, FGF-23, TNF, TWEAK, and Flt3L) are displayed separately. Linear regression analysis was performed for each biomarker, and the corresponding R^2^ and P-values are shown. (B) Volcano plot showing the association between circulating proteins measured in the UK Biobank and TAA, plotted as hazard ratio versus -Log_10_(p-value). A total of 357 proteins were positively associated and 94 proteins were negatively associated with TAA. (C) Forest plot showing the hazard ratios for the top 10 positively associated (top) and top 10 negatively associated (bottom) biomarkers in the UK Biobank dataset. (D) Forest plot showing hazard ratios for the candidate biomarkers from panel A within the UK Biobank dataset.

### FGF-23 Targets Adventitial Fibroblasts via FGFR1 and Induces Inflammatory and Matrix Remodeling Programs

To identify potential protein-protein interactions with FGF-23, we performed a network analysis using STRING^45^ (**Supplemental Figure 2A**), which highlighted interactions with FGF receptors (FGFR1–4) and the co-receptor Klotho^46^, all of which are localized to the plasma cell membrane^47^. To evaluate receptor expression in aneurysmal tissue, we analyzed our scRNA-seq dataset and found that FGFR1 was the only FGF receptor detected at meaningful levels, with expression across ECs, VSMCs, and fibroblasts, and highest levels in fibroblasts and myofibroblasts (**Figure 5A**). To further assess whether FGFR1 signaling is transcriptionally active in aneurysmal fibroblasts, we performed module score analysis of FGFR1-responsive gene sets in fibroblasts from the publicly available dataset generated by Chou et al. ^15^. This analysis revealed a modest increase in the core FGFR1 module and a significant enrichment of broader FGFR1/MAPK signaling, along with downstream inflammatory and extracellular matrix remodeling programs in aneurysmal fibroblasts compared to controls (**Figure 5B**). Consistent with these findings, inspection of the underlying module genes demonstrated that this enrichment was driven primarily by immediate-early MAPK response genes, including *EGR1*, *FOS*, *FOSB*, and *JUN* (**Supplemental Figure 2B**). In contrast, while *FGFR1* was also expressed in VSMCs, module score analysis revealed a more limited transcriptional response, with enrichment restricted to the FGFR1/MAPK program and no activation of core FGFR1 or downstream remodeling modules (**Supplemental Figure 2C**).

**Figure 5.**
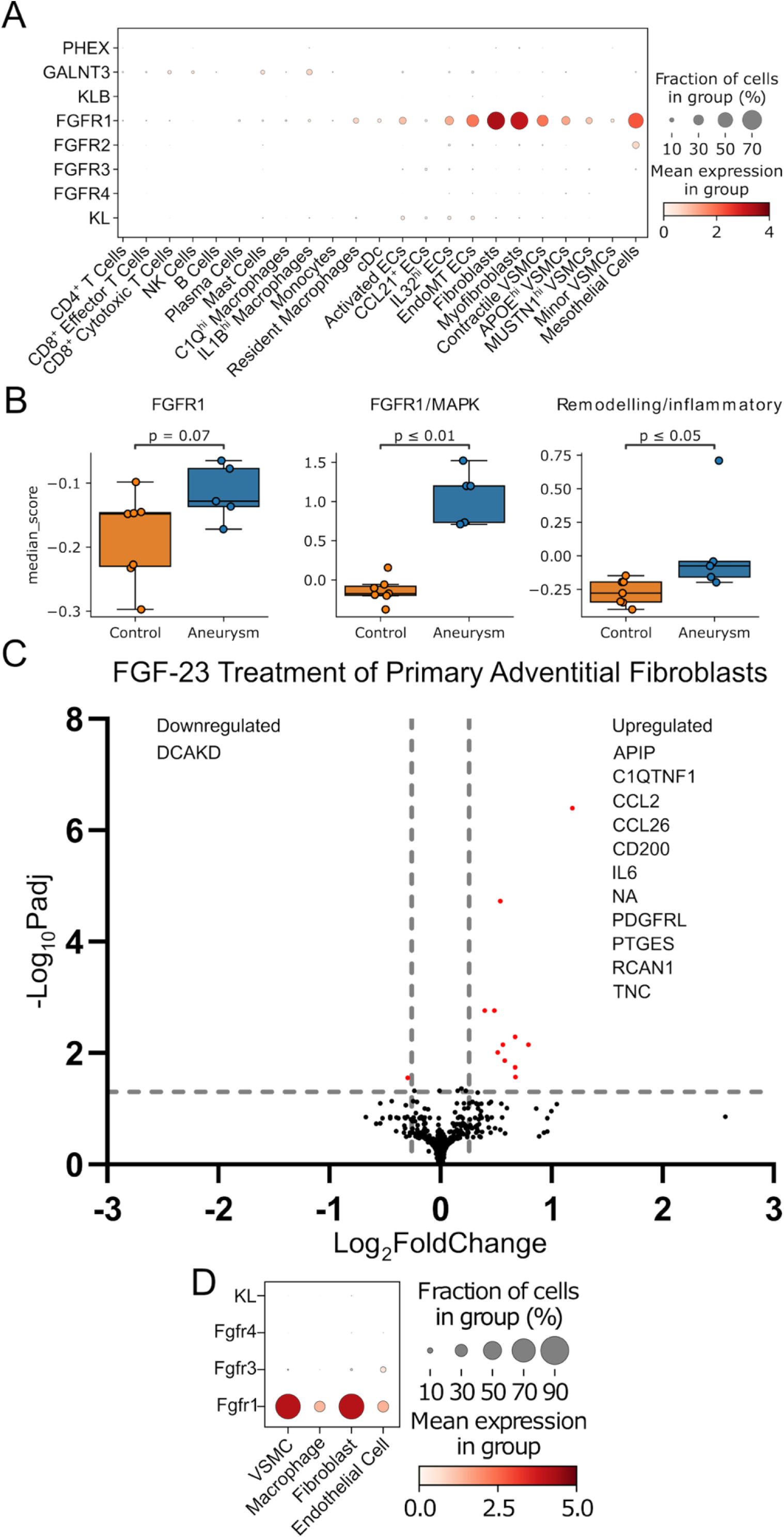
FGFR1-associated signaling is enriched in fibroblasts and induced by FGF-23. (A) Dot plot showing the expression of FGF-23-related receptors (PHEX, GALNT3, KLB, FGFR1-4, and KL) across cell populations identified by scRNA-seq in human TAA tissue. (B) Module score analysis of FGFR1 signaling-related gene set in fibroblasts from human aortic tissue, comparing control and aneurysmal samples from Chou et al dataset. Modules include primary FGFR1 response gene set, expanded FGFR1/MAPK signaling program, and a remodeling/inflammatory gene program. (C) Volcano plot showing differentially expressed genes identified with Bulk RNA-seq in primary human adventitial fibroblasts following treatment with FGF-23. All upregulated and downregulated genes are displayed. (D) Dot plot showing the expression of FGF receptors in publicly available scRNA-seq data from angiotensin II-induced TAA mouse tissue.

To assess the functional consequences of FGF-23 signaling in the principal FGFR1-expressing cell population, we performed bulk RNA-seq on primary human adventitial fibroblasts following stimulation with recombinant FGF-23 to evaluate downstream transcriptional responses. Differential expression analysis revealed a modest but coordinated transcriptional program, including upregulation of genes involved in inflammation (e.g., *IL6*, *CCL2*), extracellular matrix remodeling (e.g., *TNC*, *RCAN1*), and immunoregulatory signaling (e.g., *CD200*) (**Figure 5C**). Although effect sizes were moderate, the consistency of pathway-level changes supports a biologically meaningful response to FGF-23 signaling in adventitial fibroblasts. To determine whether this observation was consistent across species, we analyzed publicly available scRNA-seq data from Angiotensin II-induced TAA mice^24^. Similar to the human data, *Fgfr1* was the only receptor expressed at significant levels, predominantly in VSMCs and fibroblasts (**Figure 5D**). Collectively, these findings identify adventitial fibroblasts as a key cellular target of FGF-23 signaling via FGFR1 and demonstrate a conserved, fibroblast-centered pathway across human and murine aortic tissue that may contribute to TAA pathogenesis.

## Discussion

In this study, we present an integrative evaluation of the tissue microenvironment in TAA across one original and two previously published human datasets, offering novel insights while addressing key discrepancies among studies. Our analysis reveals variations between studies in cell type abundance, differentially expressed genes associated with disease pathology, and potential causal genes linked to GWAS SNPs. Despite these differences, we established a refined cell type annotation consistent across all datasets, identifying 25 distinct cell types within TAA tissue. Notably, these include five subsets of VSMCs, three subsets of fibroblasts, and five subsets of ECs. This detailed characterization lays the foundation for understanding the functional roles of these cell types in disease progression. We also demonstrated cell type-specific enrichment of key genes previously implicated in TAA, highlighting their targeted roles in disease regulation. Finally, leveraging plasma proteomics coupled with our scRNA-seq dataset, we identified the FGF23-FGFR1 signaling pathway as a potential driver of TAA formation through its activity in VSMCs and fibroblasts.

Previous single-cell studies of the human aorta have not captured the extent of cellular diversity we now describe^15,16,24,48^. Consistent with datasets that have extensively profiled macrophage subsets in health and disease, we identified several overlapping phenotypes, including C1Q^hi^, IL1B^hi^, and resident macrophages^33^. Notably, these macrophage subtypes are also observed in other vascular diseases such as atherosclerosis^32^. However, unlike atherosclerosis, we did not detect macrophages expressing high levels of lipid metabolism genes (e.g., *APOE*, *ABCA1*, *CD36*), which characterize foam cells, suggesting overlapping yet distinct inflammatory pathology. The most striking aspect of our dataset’s cell composition was the diverse phenotypes of ECs, VSMCs, and fibroblasts identified. Among these, we uncovered several potentially dysfunctional EC subpopulations, including one undergoing EndoMT—a process linked to aneurysm pathology and vascular degradation^49–51^. During EndoMT, ECs acquire mesenchymal traits such as increased migration, permeability, and expression of markers like *TAGLN* and *ACTA2*. This process, driven by TGF-β signaling and factors such as disturbed flow and inflammation, may contribute to aneurysm formation^52^. Much less is known about IL32^+^ and CCL21^+^ ECs in aneurysms. *IL32* is induced by pro-inflammatory cytokines such as IL-1β, TNF-α, and IFN-λ^53–55^, suggesting potential crosstalk between ECs and inflammatory cells like macrophages, driving an inflammatory EC phenotype. Notably, IL32 has been identified in the aortic tissue of abdominal aneurysms but not in controls, indicating a potential pathological role^56^. Interestingly, the CCL21^+^ EC also highly expressed *FABP4*, which contributes to fatty acid transport into ECs^57^, a process that may influence proliferation and angiogenesis^58^—cellular pathways potentially critical to the development of TAA.

VSMCs are key cellular components in vascular diseases, forming the majority of cells within arterial tissue and playing essential roles in maintaining structural integrity and proper function^59,60^. In recent years, significant attention has been directed toward the role of VSMCs in vascular diseases, particularly in atherosclerosis^61–68^, and to a lesser extent, in aortic aneurysms^24,48,69^. Numerous phenotypically distinct subsets of VSMCs have been identified, including pluripotent stem-like cells capable of differentiating into diverse cell types such as fibromyocytes, osteogenic cells, inflammatory cells, foam cells, and fibrochondrocytes^64^. Additionally, other yet-to-be-characterized subsets may exist, further expanding the complexity of VSMC plasticity. To add to this, here we identified distinct subsets of VSMCs, including contractile, myofibroblasts, APOE^+^, and MUSTN1^+^. While our data cannot confirm the pathophysiological relevance of any of these cellular subsets, we hypothesize that specific VSMC subsets may contribute to this process in TAA, mirroring a phenomenon observed in mouse models of abdominal aortic aneurysms^70^. We identified a previously undocumented VSMC phenotype characterized by the specific expression of *MUSTN1*. Although the physiological function of MUSTN1 remains poorly understood, it is thought to regulate myogenic and chondrogenic lineages^71^. Moreover, MUSTN1 has been shown to be secreted by VSMCs, where it promotes ECM remodeling and plays a role in fibrosis and immune cell infiltration^72^—processes that could actively drive TAA progression. Future research is essential to elucidate the specific role of MUSTN1^+^ VSMCs in disease progression, the regulatory mechanisms governing their behavior, and their impact on the tissue microenvironment.

Discrepancies in cellular alteration in single-cell studies can arise from many technical and methodological reasons. Cell type abundance is highly influenced by the location (anatomic segment) and extent (intimal vs. adventitial vs. transmural) of tissue sampling, the digestion conditions used during sample processing, and subsequent live cell isolation^73^, all of which might confer cell type bias and contribute to discrepancies across studies. VSMCs, a predominant cell type in the human aorta, play a pivotal role in aneurysm formation. In our study, VSMCs accounted for only approximately 25% of the cells, compared to the expected 60–70% in a healthy human aorta^74^. This discrepancy may result from variability in tissue isolation and processing methods, as well as differences in the type of single-cell analysis performed (scRNA-seq vs. snRNA-seq)^75^. The studies analyzed and integrated here, including our own, are relatively small in scale; however, combining datasets from diverse research sites and technologies provides a critical step toward revealing the true biological identity and cellular diversity within TAA tissue. Meta-analyses like this have already begun to emerge in other fields, including atherosclerosis^76,77^, cancer^33,78^, and infectious diseases^79^. This approach will enable the establishment of a consensus on the tissue microenvironment, as well as the identification of key cell types and their regulators that drive disease progression.

A particularly compelling aspect of our analysis is the identification of potential biomarkers and their associated cell types in TAA. Notably, our findings highlight a potential role for FGF-23-FGFR1 signaling in both VSMCs and fibroblasts. FGF-23, primarily synthesized by osteocytes and osteoblasts^80^, is well known for its role in phosphate homeostasis, where it promotes renal phosphate excretion^81,82^. Furthermore, FGF-23 has been positively associated with all-cause mortality^83^ and CVD^84^, suggesting its involvement in the pathogenesis of diverse disease states. In our cohort, we observed a marginal association between plasma FGF-23 levels and aortic diameter in TAA. Leveraging data from the UK Biobank, we validated this association by identifying elevated FGF-23 levels in patients with TAA compared to controls. Linking this to our scRNA-seq data, we observed that *FGFR1* was the only receptor for FGF-23 expressed at significant levels in both human and mouse aneurysmal tissue, with highest expression in fibroblasts and VSMCs. To assess the functional consequences of this signaling axis, we performed bulk RNA-seq in primary human adventitial fibroblasts following FGF-23 stimulation, which revealed a coordinated transcriptional program characterized by induction of inflammatory and ECM remodeling pathways. These processes are well-established contributors to aneurysm pathogenesis, including cytokine-mediated vascular inflammation^85^ and dysregulation of the ECM^86^, suggesting that FGF-23 signaling may reinforce fibroblast-driven remodeling programs within the aortic wall. Together, these findings support a model in which circulating FGF-23 reflects and may contribute to pathogenic stromal signaling networks underlying TAA. Accordingly, FGF-23 may serve as a circulating plasma biomarker of aneurysm burden, while also highlighting a potentially significant and druggable signaling pathway and the key cell types underlying TAA pathobiology. Importantly, our study was not designed to identify biomarkers for population-level TAA screening or prediction of acute aortic events such as dissection of rupture; longitudinal validation studies will be needed to determine whether FGF-23 measurements may be useful in these settings.

In conclusion, we employed an integrative approach to characterize the cellular diversity in human TAA, with a particular focus on stromal cell populations that play a pivotal role in thoracic aortic disease. Furthermore, we identified a novel signaling pathway that may serve as both a circulating biomarker of disease burden and a contributing factor in disease progression. To fully establish the clinical utility of FGF-23 as a biomarker, further validation is required. Additionally, mechanistic studies in murine models are essential to elucidate the molecular mechanisms underlying the role of FGF-23 in disease pathology, as well as to investigate the functions of the refined cell types identified in our analysis.

### Limitations

This study has several important limitations. First, we lacked appropriate control tissue for both the single-cell RNA sequencing and plasma proteomic analyses, which restricted our ability to directly compare cellular and molecular features with those observed in third-party datasets. Future studies incorporating matched healthy aortic tissue will be critical for validating and contextualizing our findings. Additionally, the lack of precise anatomical annotation of aortic specimens complicates cross-study comparisons. Consequently, some of the observed discrepancies between studies may arise from subtle differences in tissue sampling location. Second, our dataset included one individual with Marfan syndrome and one with Loeys-Dietz syndrome. While these rare syndromic cases were deliberately included to broaden the relevance and utility of our dataset, the cellular mechanisms distinguishing syndromic from non-syndromic TAA remain incompletely defined. The inclusion of these samples provides a valuable resource for future investigations into the shared and distinct pathways contributing to TAA pathogenesis across genetic backgrounds. Third, although our data suggests a potential association between circulating FGF-23 levels and TAA, these findings are correlative and primarily descriptive. Establishing a causal role for FGF-23 will require the development of *in vivo* models to dissect its mechanistic contribution to aneurysm initiation and progression. Finally, our targeted plasma proteomic analysis was performed on a small number of individuals sampled at the most severe stage of disease. To more fully capture the temporal dynamics of potential protein biomarkers, future studies should incorporate larger, longitudinal cohorts with serial sampling across the course of disease development. Such efforts would facilitate the identification of stage-specific biomarkers and provide insight into the molecular transitions that characterize TAA progression. Although it is not surprising that our in-house cohort was predominantly male—given the convenience sampling approach and known sex skew of TAA—the other studies analyzed here exhibited greater sex balance. Expanding our cohort size will enable a more thorough analysis of sex-specific differences, as well as variations across diverse demographic and clinical parameters.

### Competency in Medical Knowledge

Thoracic aortic aneurysm is managed primarily through imaging surveillance and diameter-based thresholds for intervention, yet the cellular programs that underlie disease progression and the circulating factors that reflect disease burden remain incompletely defined. In this study, integrative single-cell transcriptomic and plasma proteomic analyses identified substantial stromal and immune heterogeneity in human thoracic aortic aneurysm tissue and nominated FGF-23 as a candidate circulating biomarker associated with aneurysm burden and with the presence of thoracic aortic aneurysm in the UK Biobank. We further show that FGFR1, a receptor for FGF-23, is most prominently expressed in fibroblasts and subsets of vascular smooth muscle cells, and that FGF-23 induces inflammatory and extracellular matrix remodeling programs in primary human adventitial fibroblasts. These findings refine current understanding of thoracic aortic aneurysm biology and highlight the FGF-23–FGFR1 axis as a candidate stromal signaling pathway relevant to disease progression.

### Translational Outlook

This study identifies FGF-23 as a candidate biomarker of thoracic aortic aneurysm burden and highlights the FGF-23–FGFR1 axis as a candidate therapeutic target in disease progression. Further studies are needed to determine whether FGF-23 adds value beyond imaging for identifying disease presence, severity, or progression, and whether it may be useful for longitudinal monitoring. In parallel, mechanistic studies in experimental models will be necessary to determine whether targeting FGF-23–FGFR1 signaling can attenuate fibroblast-centered inflammatory and extracellular matrix remodeling programs in thoracic aortic aneurysm.

### Sources of Funding

A.C.B is supported by a Career Development Award from the Marfan Foundation. E.J.G is supported by an American Heart Association Postdoctoral Fellowship (25POST1369638).

## Abbreviations and Acronyms

TAA: Thoracic Aortic Aneurysm
ECM: Extracelluar Matrix
VSMC: Vascular Smooth Muscle Cell
scRNA-seq: Single-Cell RNA-Sequencing
EC: Endothelial Cell
CMR: Cardiac Magnetic Resonance
MRA: Magnetic Resonance Angiography
CTA: Contrast Tomography Angiography
GEX: Gene Expression
PCA: Principal Component Analysis
UMAP: Uniform Manifold Approximation and Projection
PCR: Polymerase Chain Reaction
DE: Differentially Expressed
snRNA-seq: Single-Nuclei RNA-Sequencing

