## Supplemental Appendix for "Integrative Omics Identifies Candidate Plasma Biomarkers and Cellular Targets Associated with Thoracic Aortic Aneurysm"

**Supplemental Figure 1.** Reanalysis of single-cell transcriptomic data after excluding individual subjects.

**Supplemental Figure 2.** FGFR1-associated signaling and gene module expression across cell types.

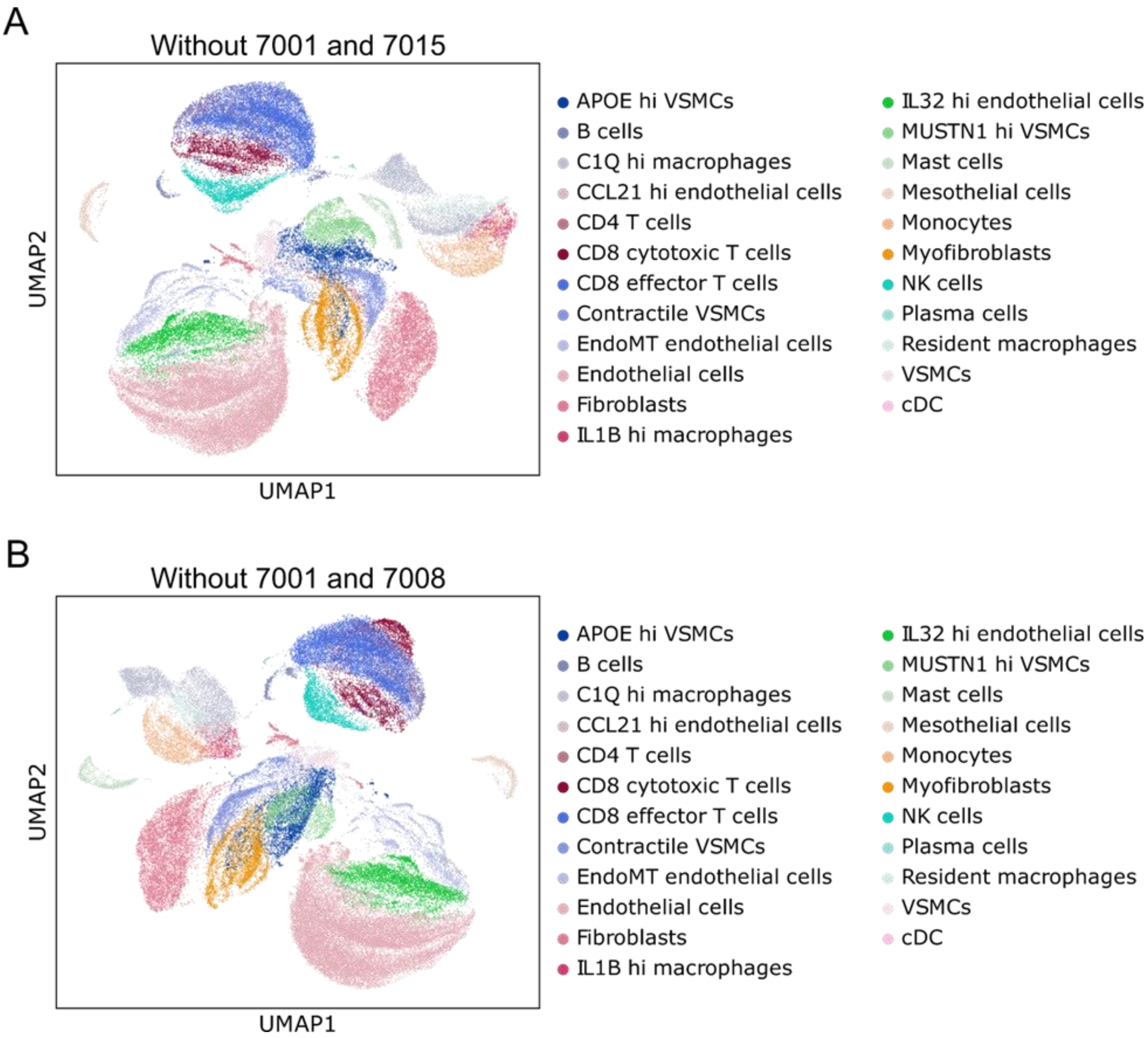


**Supplemental Figure 1. Reanalysis of single-cell transcriptomic data after excluding individual subjects.** (A) UMAP visualization of cell populations obtained after removing subjects 7001 and 7015. (B) UMAP visualization of cell populations after removing subjects 7001 and 7008.

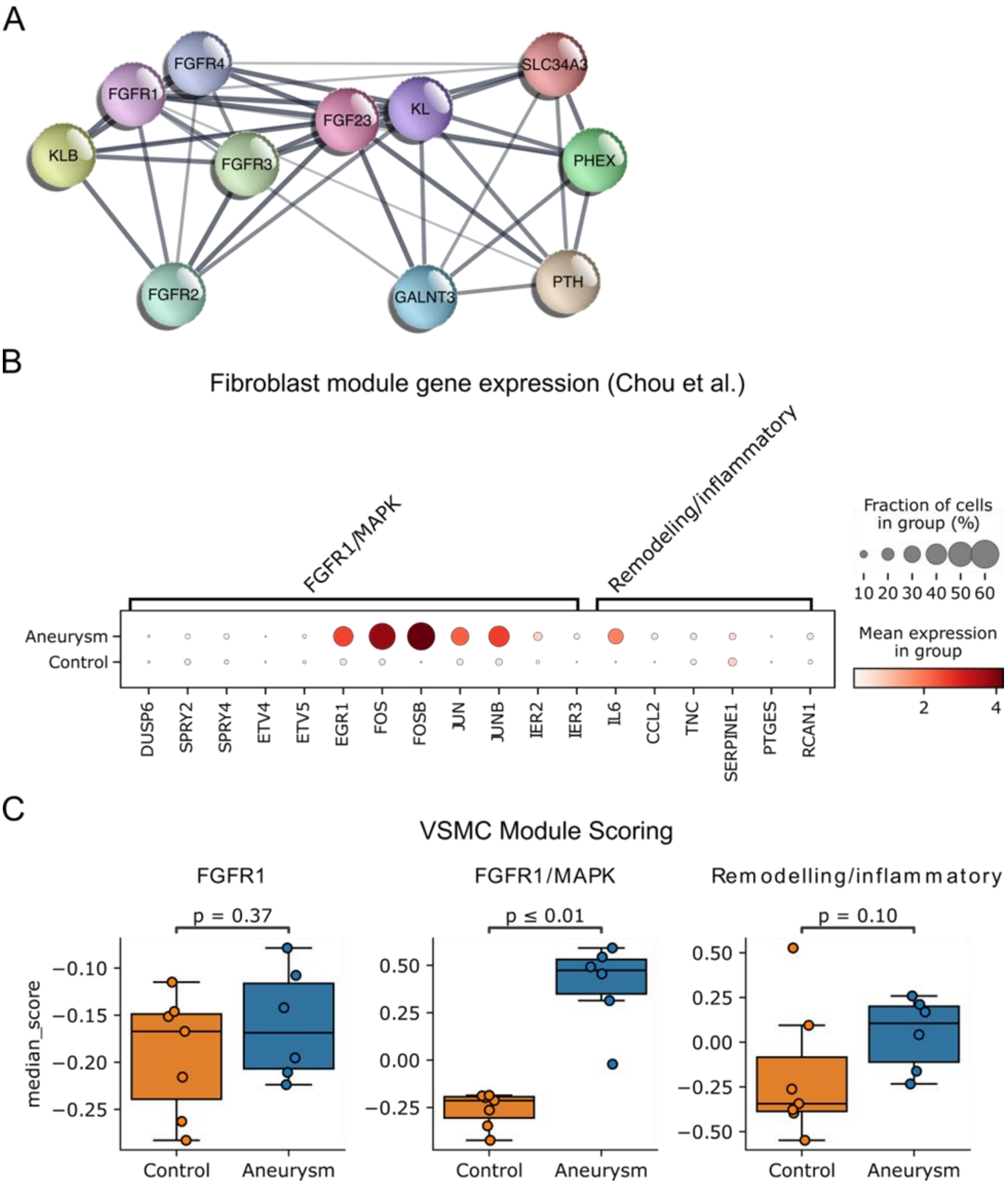


**Supplemental Figure 2. FGFR1-associated signaling and gene module expression across cell types.** (A) STRING network analysis of FGF-23 and predicted protein–protein interactions. (B) Dot plot showing expression of genes included in FGFR1-associated transcriptional modules in fibroblasts from dataset by Chou et al. (C) Module score analysis of FGFR1-associated gene sets in vascular smooth muscle cells (VSMCs) from the Chou et al. dataset.
